# PARP1-mediated PARylation of SPRTN restrains its DNA-protein crosslink repair activity

**DOI:** 10.64898/2026.11.04.764412

**Authors:** Ayesha Rahman, Veronika Quevedo, Martin Garcia, Andrew Rhiner, Quincee Simonson, Estella Perez, Jaime Lopez-Mosqueda

## Abstract

SPRTN is a DNA-dependent metalloprotease that resolves DNA-protein crosslinks, but how its activity is restrained on chromatin remains unclear. Here we show that SPRTN is PARylated in a PARP1- and HPF1-dependent manner. Mass spectrometry datasets identify seven modified residues in the C-terminal region, and substituting them (SPRTN-PARless) strongly reduces PARylation. The Nudix homology domain is required for SPRTN to interact with PARP1 and for efficient PARylation. SPRTN-PARless accumulates faster and to higher levels at laser-induced DNA damage than wild-type SPRTN. PARylation reduces single-stranded DNA binding of both cell-derived and purified SPRTN, and PARG treatment reverses this effect for cell-derived protein. SPRTN-PARless shows increased autocleavage and accelerates clearance of camptothecin-induced TOP1 cleavage complexes. These findings suggest that PARylation acts as negative feedback on SPRTN after its PAR-dependent recruitment, restraining its DNA engagement and proteolytic activity.

## INTRODUCTION

Genome integrity is continually challenged by DNA lesions arising from both endogenous and exogenous sources, and cells rely on a coordinated DNA damage response (DDR) to detect, signal, and repair this damage^1,2^. A particularly toxic class of DNA lesions are DNA-protein crosslinks (DPCs), in which proteins become covalently trapped on DNA, physically blocking replication and transcription machinery^3^. DPCs arise from a variety of sources, including reactive aldehydes such as formaldehyde, chemotherapeutic agents such as topoisomerase poisons, and covalent enzymatic intermediates that become trapped on DNA, exemplified by topoisomerase I and II cleavage complexes (TOP1cc and TOP2cc)^3,4^. Left unresolved, DPCs promote genomic instability, premature aging, and cancer predisposition.

Several pathways converge to resolve DPCs, including nucleotide excision repair (NER) and homologous recombination (HR)-associated mechanisms for smaller crosslinked peptides, and dedicated proteolytic pathways for bulkier protein adducts. SPRTN was identified as the first mammalian metalloprotease capable of proteolytically resolving DPCs and is now recognized as a central component of DPC repair^5–7^. In addition to SPRTN, the 26S proteasome and other proteases, including FAM111A and the SPRT-domain protein GCNA/ACRC, have been implicated in the processing of specific DNA-protein adducts^8–10^. SPRTN can process a broad range of chromatin-associated substrates, including histones, DNA topoisomerases, and PARP1^3,11^. SPRTN’s activity is essential in mice and biallelic hypomorphic mutations in human SPRTN cause Ruijs-Aalfs syndrome, a disorder characterized by genome instability, segmental progeria, and early-onset hepatocellular carcinoma, underscoring the physiological importance of proper SPRTN function^12^.

Because SPRTN’s protease activity shows only limited substrate specificity beyond a general preference for DNA-proximal, disordered protein regions, its recruitment and activation must be tightly restrained in space and time to avoid indiscriminate degradation of chromatin-bound proteins. Multiple layers of regulation have been proposed to control SPRTN activity and localization, including cell-cycle-dependent phosphorylation, monoubiquitination that autoinhibits its protease domain, and recruitment via its SHP and PIP-box motifs to p97/VCP and PCNA at sites of replication-associated damage ^13–16^. Recent studies have further revealed that ubiquitin can directly stimulate SPRTN proteolytic activity, providing a mechanism by which ubiquitinated DPCs may selectively activate SPRTN at sites of damage^17,18^. These findings raise the broader question of how opposing regulatory signals are integrated to control SPRTN recruitment and proteolysis in the context of DNA damage.

Poly(ADP-ribosyl)ation (PARylation) is a rapid and reversible signal for coordinating DDR factor recruitment at diverse lesion types. PARP1, the founding member of the PARP family, is activated upon binding to damaged DNA and, together with the histone PARylation factor HPF1, catalyzes DNA damage-induced PARylation of histones and other chromatin-associated proteins^19–22^. PAR polymers are subsequently removed by PARG and other dePARylating enzymes such as ARH3, restoring the local chromatin environment^23,24^. Recent work has begun to implicate this signaling axis directly in DPC repair. Hurley et al. recently reported that PARP1 and PARP2 sense DPCs and modify them with PAR, and identified a Nudix homology domain (NHD) within SPRTN that mediates direct, non-covalent PAR binding^25^, distinguishing it from the canonical zinc-binding domain previously proposed for this region^26^. Loss of PARP1/2 activity or mutation of the SPRTN NHD impaired DPC repair, and single-molecule analysis showed that SPRTN is recruited to DPCs inefficiently on its own, but binds following PARP1-dependent PARylation of chromatin in the presence of NAD+, establishing PAR as a recruitment signal for SPRTN at sites of DPC damage^25^.

While this work establishes that SPRTN is recruited to DPCs via PAR sensing, whether SPRTN itself is a direct substrate for PARylation, and how such a modification might regulate its activity once recruited, has not been examined. Here we show that SPRTN is directly PARylated by PARP1 in an HPF1-dependent manner, and we identified seven PARylation sites on SPRTN by mining published proteome-wide mass spectrometry datasets. Using a PARylation-deficient mutant (SPRTN^PARless^), we identify PARylation as a key mechanism that uncouples SPRTN recruitment from proteolytic activation. Although PARP1/2 activity is required for efficient SPRTN recruitment to DNA damage sites, SPRTN PARylation limits its accumulation on damaged chromatin and suppresses its DNA-binding. Consistent with this inhibitory role, SPRTN^PARless^ accumulates more at microirradiation-induced DNA damage sites than SPRTN^WT^ and exhibits increased self-cleavage activity in cells, resulting in accelerated resolution of TOP1ccs. PARG-mediated PAR removal reverses the inhibition of SPRTN DNA binding, while PARG inhibition reduces chromatin-associated SPRTN, further supporting PAR-dependent regulation of SPRTN association with damaged chromatin. Together, these findings support a model in which a reversible PARylation cycle fine-tunes SPRTN activity after recruitment, ensuring that DPC proteolysis is activated only at the appropriate stage of repair.

## RESULTS

### SPRTN is modified with poly(ADP-ribosyl)ation

During experiments involving transient expression of SPRTN, we repeatedly observed a high-molecular-weight species associated with the protein. Given the requirement for PARP1/2 activity in SPRTN function at sites of DNA damage, we hypothesized that these species might represent PARylated forms of SPRTN. To test this possibility, GFP-SPRTN^WT^ was transiently expressed in HEK293T cells and immunoprecipitated with GFP trap beads. Immunoprecipitates revealed both full-length SPRTN and a series of higher-molecular-weight species that were absent from GFP controls (**Figure 1A**). Notably, treatment with the PARP inhibitor Olaparib markedly reduced both the high-molecular-weight SPRTN species and the associated PAR signal (**Figure 1B**), indicating that SPRTN undergoes PARP-dependent modification in cells.

**Figure 1:**
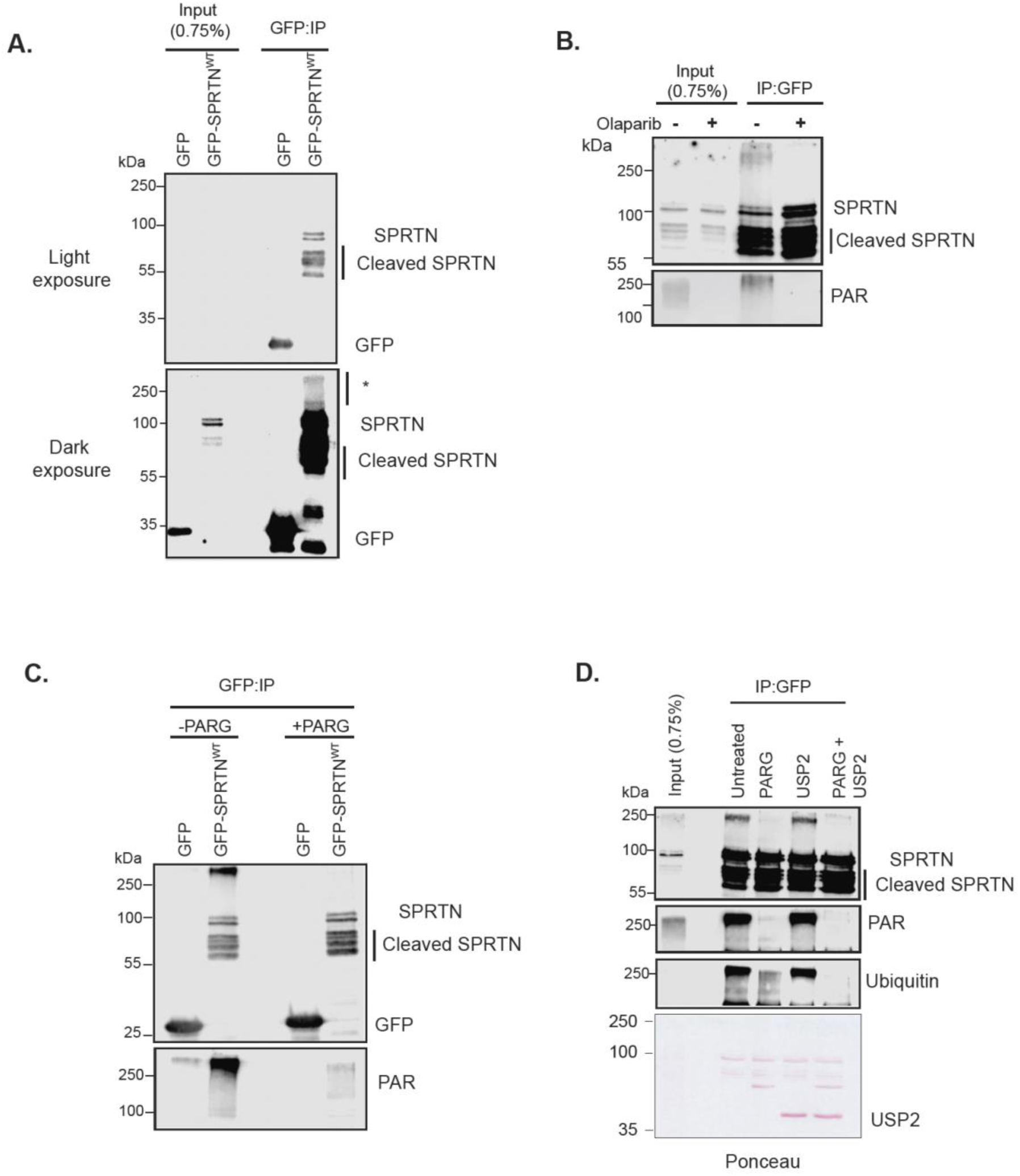
SPRTN is modified with poly(ADP-ribosyl)ation. **A.** Immunoprecipitation (IP) of GFP and GFP-SPRTN^WT^. High-molecular-weight SPRTN species were observed under dark exposure. **B.** Effect of PARP inhibition. Cells were either untreated or treated for 16 hours with 20 μM Olaparib prior to IP. **C.** PARG treatment. Post-IP samples were split into two fractions and treated with or without PARG for 30 min. **D.** Combined PARG and USP2 treatment. Post-IP samples were split into four fractions and subjected to no treatment, PARG treatment (1 hour), 2 μM USP2 treatment (1 hour), or combined PARG+USP2 treatment (1 hour). All experiments (A-D) were conducted after overexpressing GFP and GFP-SPRTN^WT^ by transient transfection in HEK293T cells as indicated. 20 μM PARGi was added (16 hours) for B-D.

To confirm directly that SPRTN is post-translationally modified with PAR, we treated GFP-SPRTN^WT^ immunoprecipitates with poly-(ADP-ribose)-glycohydrolase (PARG). PARG treatment collapsed the high-molecular-weight GFP-SPRTN^WT^ species into discrete lower-molecular-weight species and reduced the associated PAR signal, whereas GFP-only controls lacked these high-molecular-weight species (**Figure 1C**). These results demonstrate that the high-molecular-weight SPRTN species are modified with poly(ADP-ribose) polymers. Because SPRTN has also been reported to be ubiquitinated^27–29^, we asked whether the high-molecular-weight species on GFP-SPRTN^WT^ reflects PARylation, ubiquitination, or both. To distinguish PARylation from ubiquitination, we treated GFP-SPRTN immunoprecipitates with PARG, the deubiquitinase USP2, or both. PARG removed the PAR signal and the high-molecular-weight SPRTN species, whereas USP2 reduced the ubiquitin signal but left a prominent high-molecular-weight species intact. This residual species was lost only upon PARG treatment, whether alone or in combination with USP2 (**Figure 1D**), indicating that PARylation, rather than ubiquitination, is the principal contributor to the high-molecular-weight SPRTN species.

### SPRTN is PARylated by PARP1 in an HPF1-dependent manner

To test whether PARP1 can modify SPRTN directly, we reconstituted the reaction *in vitro* using recombinant PARP1, SPRTN, NAD+, HPF1, and nucleosomes. Incubation of SPRTN with PARP1 under these conditions generated a high-molecular-weight SPRTN species that was absent when PARP1 was omitted (**Figure 2A**), indicating that PARP1 can modify SPRTN *in vitro*. Because Olaparib inhibits both PARP1 and PARP2, we next examined the contribution of PARP1 specifically in cells. We immunoprecipitated GFP-SPRTN from control and PARP1-depleted cells and found that SPRTN PARylation was reduced upon PARP1 knockdown (**Figure 2B**), indicating that SPRTN PARylation in cells depends on PARP1. PARP1 modifies serine residues in complex with its cofactor HPF1, which also limits the length of the PAR chains that PARP1 synthesizes^30^. Consistent with this, inclusion of HPF1 in our *in vitro* reactions reduces the high-molecular-weight SPRTN species (**Figure 2A**). We therefore asked whether SPRTN PARylation in cells also requires HPF1. Immunoprecipitation of GFP-SPRTN from control and HPF1-depleted cells reveals a loss of SPRTN PARylation in the absence of HPF1 (**Figure 2C**), indicating that HPF1 is required for SPRTN PARylation in cells. Together, these data indicate that SPRTN is PARylated in a PARP1-and HPF1-dependent manner.

**Figure 2:**
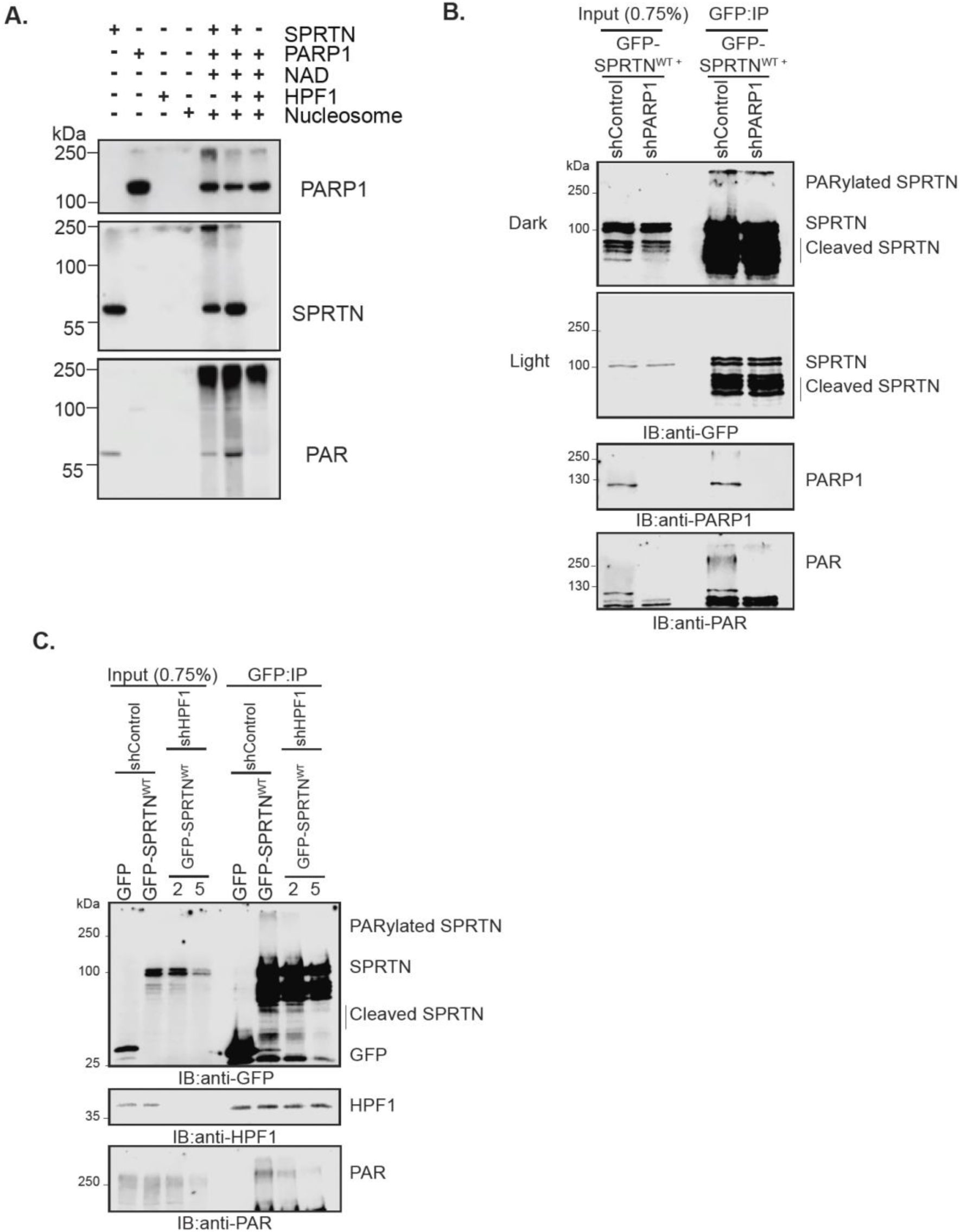
SPRTN is PARylated by PARP1 in an HPF1-dependent manner. **A.** In vitro PARylation of SPRTN by PARP1. 1.25 pmol PARP1 and 25 pmol HPF1 were used to in vitro PARylate purified full length SPRTN (0.5 µM). NAD+ (500 µM) served as a substrate for PAR production. **B.** Immunoprecipitation of GFP-SPRTN^WT^ from HEK293T shControl and B. shPARP1 or **C.** two different shHPF1 cell lines. GFP-SPRTN^WT^ was expressed by transient transfection in HEK293T cells as indicated. 5 µM PARGi was added (16 hours) for B-C.

### SPRTN is PARylated at seven mapped residues clustered in its C-terminus

To identify the residues on SPRTN that are PARylated, we mined published proteome-wide ADP-ribosylation datasets for SPRTN modification sites. Two independent datasets identified overlapping PARylation sites on SPRTN^31,32^, comprising six serine residues (S254, S275, S373, S382, S383, S429) and one lysine residue (K376), distributed across and adjacent to the SHP and PIP boxes and near the C-terminal UBZ domain (**Figure 3A**). The predominance of serine residues is consistent with the HPF1 dependence shown above, as HPF1 redirects PARP1 activity toward serine. We validated two of these sites, S383 and S429, by manual interpretation of the underlying MS/MS spectrum, which showed diagnostic c- and z-ions localizing the modification to S383 and S429 within the ISLRNSSK and EQIKSSGNDPK peptides, respectively (**Figure 3B**).

**Figure 3:**
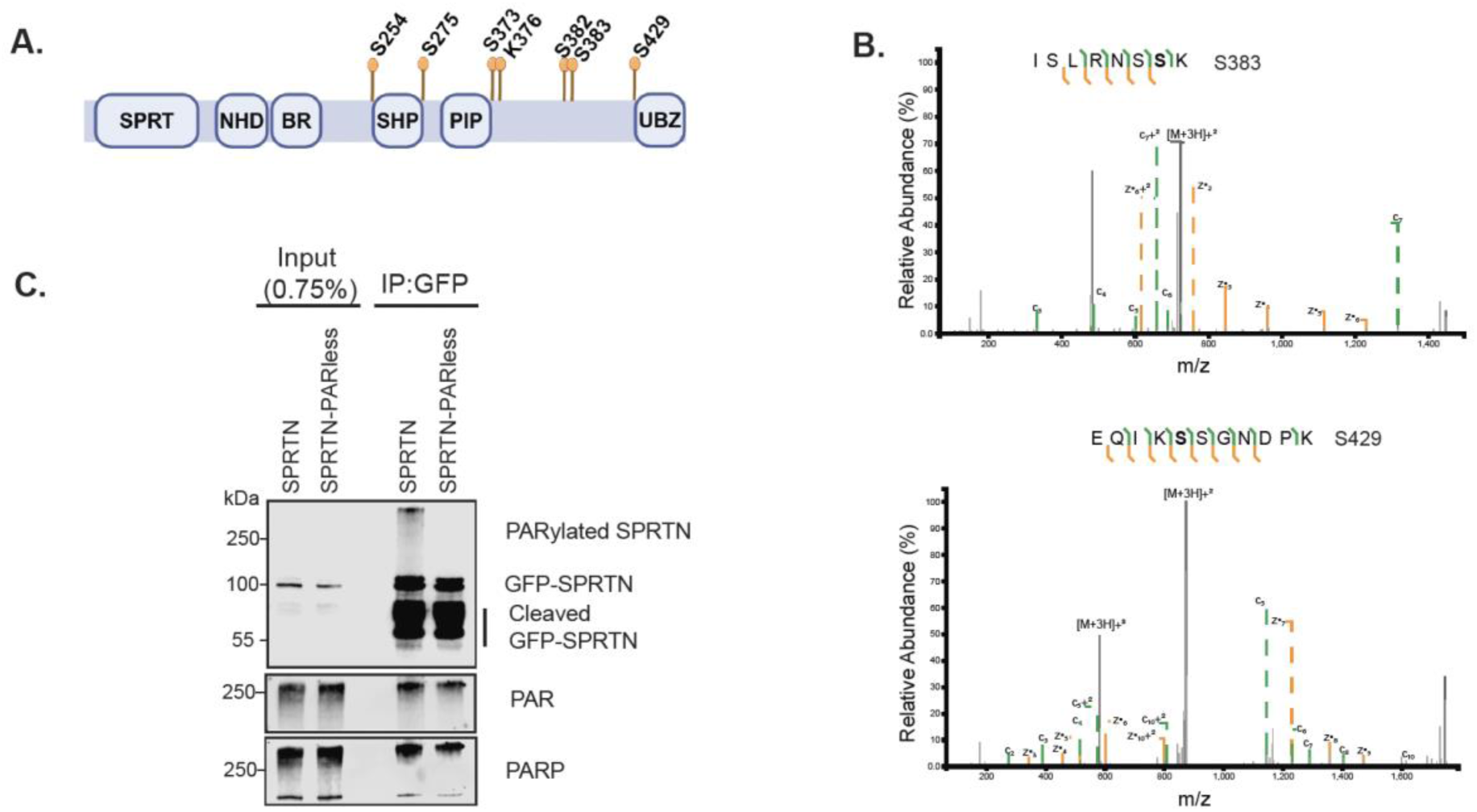
PARylation sites of SPRTN. **A.** Schematic representation of full-length SPRTN showing 7 PARylation sites identified by mass spectrometry^31–32^. **B.** Example of mass spectra of two PARylation sites (S383, S429). For visualization, only relevant ions are labeled. Peak positions and intensities are kept unaltered as the original spectra. **C.** GFP tagged SPRTN^WT^ and SPRTN^PARless^ were overexpressed and immunoprecipitated from HEK293T cells. GFP-SPRTN^PARless^ lacks high mw PARylated SPRTN species.

Because the mapped sites cluster toward the C-terminus, we tested whether this region is required for PARylation. A SPRTN mutant lacking the C-terminus (SPRTNΔC) was not PARylated (**Figure S1**), consistent with the C-terminus being the major site of modification. We then generated a SPRTN mutant in which the six identified serines residues were substituted with alanine and the lysine with arginine (S254A, S275A, S373A, K376R, S382A, S383A, S429A), which we termed SPRTN^PARless^. To confirm that this mutant is indeed refractory to PARylation, we immunoprecipitated GFP-SPRTN^WT^ or GFP-SPRTN^PARless^ from HEK293T cells and probed for PAR. While SPRTN^WT^ showed the characteristic high-molecular-weight PARylated species, this signal was strongly reduced on SPRTN^PARless^ despite comparable levels of immunoprecipitated protein (**Figure 3C**), confirming that these seven residues account for the majority of SPRTN PARylation in cells. Together, these data establish that SPRTN is PARylated predominantly at a cluster of C-terminal residues and that SPRTN^PARless^ provides a tool to define the functional consequences of this modification.

### SPRTN PARylation restrains its accumulation at sites of DNA damage

We next asked how SPRTN PARylation relates to its recruitment to DNA damage. We generated localized damage by UV-laser microirradiation and compared the kinetics of GFP-PARP1, GFP-SPRTN^WT^ and GFP-SPRTN^PARless^ by live-cell imaging (**Figure 4A**). GFP-PARP1 was recruited within seconds and rapidly dissipated. In contrast, GFP-SPRTN^WT^ accumulated after PARP1 and was retained at damage tracks for the duration of the time course. Consistent with our previous finding, SPRTN recruitment depended on PARP1/2 activity^25^. In U2OS PARP1/2 double-knockout cells^33^, GFP-SPRTN^WT^ showed minimal accumulation, comparable to background. Notably, GFP-SPRTN^PARless^ accumulated earlier and to a higher intensity than SPRTN^WT^ (**Figure 4A and 4B**). Thus, SPRTN’s own PARylation is not required for its recruitment, but instead limits the speed and extent of its accumulation at damage sites. We therefore asked how SPRTN engages PARP1. The SPRTN NHD contains the PAR-binding domain, and its mutations disrupt PAR binding, damage site recruitment and DPC repair^25^. GFP-SPRTN^WT^ co-immunoprecipitated PARP1 and carried substantial PAR modification, whereas the GFP-SPRTN^NHD^ was PARylated less efficiently and failed to interact with PARP1 despite comparable expression (**Figure 4C**). These results suggest that SPRTN is first recruited to PARP1 through PAR binding via its NHD and is then PARylated by PARP1. Finally, treating cells with H2O2 to trigger global PARP1 activation^34^, increased cellular PAR and markedly increased PARylated GFP-SPRTN^WT^, as detected by anti-GFP and anti-PAR immunoblotting (**Figure 4D**), confirming that SPRTN PARylation is inducible by DNA damage. Together, these data support a model in which PARP1 activity recruits SPRTN, and PARP1, then PARylates SPRTN, which in turn tempers SPRTN’s accumulation at damage sites.

**Figure 4:**
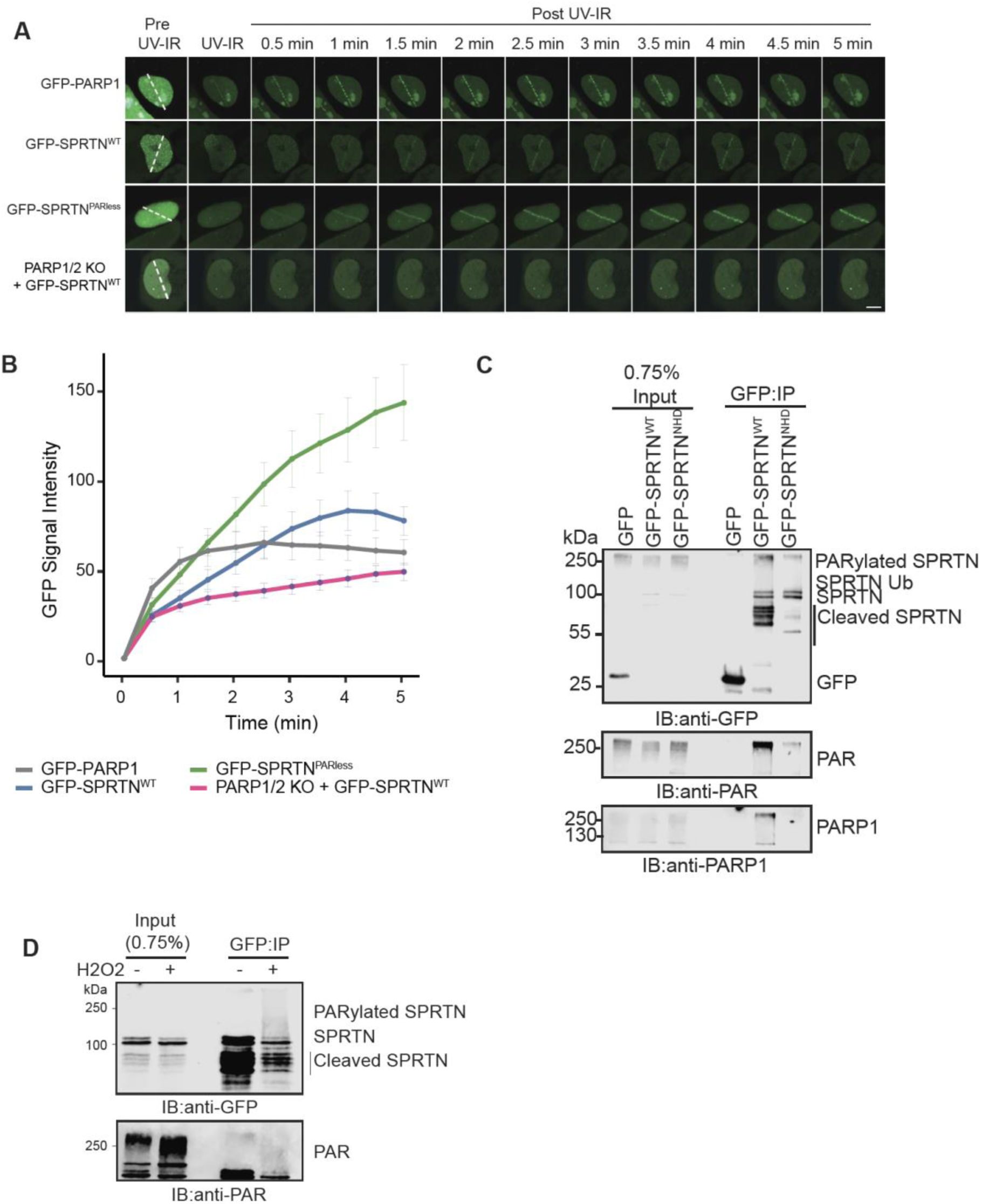
SPRTN PARylation restrains its accumulation at sites of DNA damage. **A.** UV laser microirradiation (UV-IR) was performed in U2OS cells expressing GFP-PARP1, GFP-SPRTN^WT^ or GFP-SPRTN^PARless^, and in U2OS PARP1/2 KO cells expressing GFP-SPRTN^WT^. Representative confocal images are shown before (Pre-UV-IR), during (UV-IR) and after damage induction (Post-UV-IR). The white dotted line indicates the UV damage track. Recruitment of GFP tagged proteins to the damage line was monitored for up to 5 minutes. Approximately 20-30 nuclei were damaged for each condition. U2OS PARP1/2 KO cells were treated with 20 μM Olaparib. Scale bar represents 10 μm. **B.** Quantification of the UV-IR time course experiment showing recruitment kinetics of GFP-PARP1, GFP-SPRTN^WT^ and GFP-SPRTN^PARless^ to the UV damage line. The x-axis indicates time (min) and the y-axis represents normalized GFP fluorescence intensity. Error bars represent mean ± SEM from ∼20-30 nuclei for each condition. **C.** Immunoprecipitation of GFP, GFP-SPRTN^WT^ and GFP-SPRTN^NHD^ from HEK293T to assess SPRTN-PARP1 interaction and PARylation. **D.** GFP-SPRTN^WT^ is immunoprecipitated from HEK293T cells that were either untreated or H2O2 (5 mM for 15 minutes) to observe SPRTN PARylation.

### PARylation suppresses SPRTN accumulation at damage sites and reduces SPRTN-DNA binding

SPRTN’s own PARylation restrains its accumulation at damage sites under basal conditions (**Figures 4A-B**). We therefore asked whether globally elevating PARylation with H2O2 would further suppress SPRTN accumulation at laser-induced damage. We pretreated cells with H2O2 minutes prior to UV-laser microirradiation and monitored accumulation of GFP-SPRTN^WT^, GFP-SPRTN^PARless^, or GFP-SPRTN^WT^ in PARP1/2 knockout cells (**Figure 5A**). GFP-SPRTN^WT^ accumulation was substantially blunted relative to untreated cells, plateauing at roughly one-third the intensity reached by GFP-SPRTN^PARless^ or by GFP-SPRTN^WT^ in PARP1/2 knockout cells, both of which reached comparably high plateau intensities (**Figure 5B**). Because this H2O2-induced suppression of accumulation was relieved both by removing SPRTN’s PARylation sites and by removing PARP1/2 itself, we conclude that H2O2-induced PARP1 activation inhibits SPRTN accumulation at damage specifically through SPRTN’s own PARylation, rather than through a PARP1/2-independent effect of H2O2.

**Figure 5:**
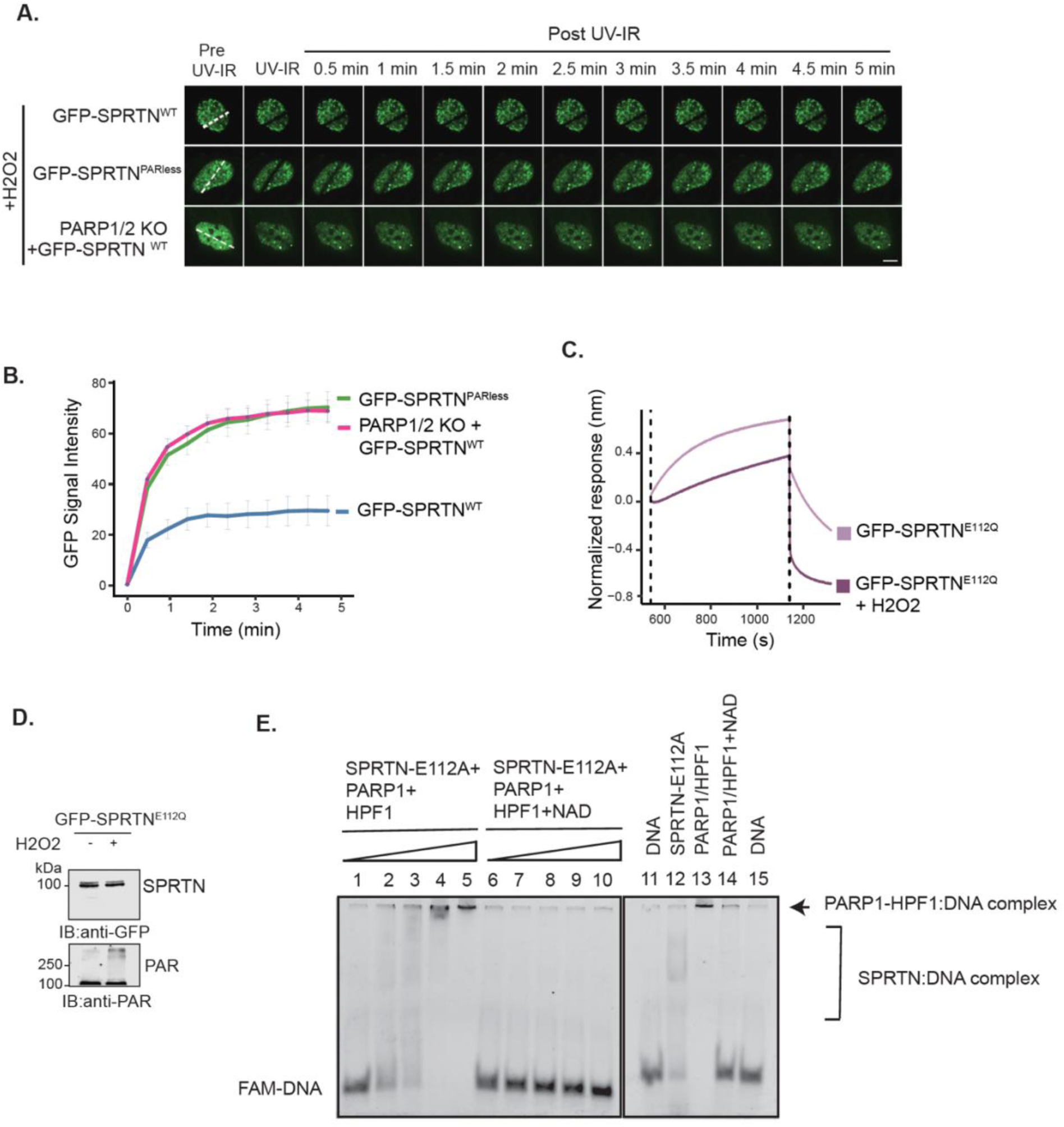
PARylation suppresses SPRTN accumulation at damage sites and reduces SPRTN-DNA binding. **A.** UV laser mediated microirradiation (UV-IR) was performed in U2OS cells expressing GFP-SPRTN^WT^ and GFP-SPRTN^PARless^. Representative confocal images are shown before (Pre-UV-IR), during (UV-IR) and after damage induction (Post-UV-IR). The white dotted line indicates the UV damage track. Recruitment of GFP tagged proteins to the damage track was monitored for up to 5 minutes. Approximately 30-50 nuclei were damaged for each condition. Cells were treated with H2O2 (5 mM) prior to UV damage. Scale bar represents 10 μm. **B.** Quantification of the UV-IR time course experiment showing recruitment kinetics of GFP-SPRTN^WT^ and GFP-SPRTN^PARless^ to the UV damage track following H2O2 treatment. The x-axis represents time (min) and the y-axis represents normalized GFP fluorescence intensity. Error bars represent mean ± SEM from ∼30-50 nuclei for each condition. **C.** Biolayer interferometry (BLI) assay. Biotinylated 20-mer ssDNA was immobilized on streptavidin biosensor and whole-cell extracts from HEK293T overexpressing GFP-SPRTN^E112Q^, treated with or without H2O2 (5 mM, 15 minutes) were used to detect SPRTN-DNA binding. DNA association and dissociation were monitored for 10 minutes and 3 minutes respectively. The x-axis represents time (s) and y-axis represents normalized binding response (nm). **D.** Western blot showing GFP-SPRTN^E112Q^ expression and PARylation levels from the cellular extracts used in Figure 4C. **E.** Electrophoretic mobility shift assay (EMSA) to test SPRTN-DNA binding. Purified SPRTN^E112Q^ of different concentrations (0.125 nM, 0.25 nM, 0.5 nM and 1 uM) were used as indicated, along with 1 uM PARP1 or HPF1 and 2.5 nM FAM labeled ssDNA.

We next examined whether PARylation affects SPRTN DNA binding. To test this, we used doxycycline-inducible stable cell lines expressing Flag-HA-Flag-SPRTN^WT^ (Flag-SPRTN^WT^) (**Figure S2**). Cells were treated with either a PARG inhibitor (PARGi) or the PARP inhibitor Olaparib. Using chromatin fractionation, we observed that under untreated conditions, Flag-SPRTN^WT^ was readily detected in the chromatin-bound fraction. However, upon PARG inhibition, chromatin bound SPRTN levels were reduced, whereas treatment with Olaparib did not noticeably affect SPRTN chromatin association (**Figure S2**).

To ask whether PARylation affects SPRTN’s affinity for DNA, we performed biolayer interferometry (BLI) with immobilized single-stranded DNA and extracts from H2O2-treated or untreated cells expressing catalytically inactive GFP-SPRTN^E112Q^, which avoids confounding autocleavage. H2O2 pretreatment reduced both the association and the apparent affinity of GFP-SPRTN^E112Q^ for the DNA-coated sensor, consistent with cellular PARylation antagonizing SPRTN-DNA binding (**Figure 5C**). As a control for the extent of PARylation induced under these conditions, we immunoblotted a sample of the cell extracts used from the same cells used in Figure 5C and confirmed increased PAR signal relative to untreated cells (**Figure 5D**).

As an additional cellular confirmation of this result, we expressed GFP-SPRTN^E112Q^, GFP-SPRTN^PARless-E112Q^, and GFP-SPRTN^NHD-E112Q^ in HEK293T cells, immobilized them on GFP-Trap beads, and incubated the beads with a Cy5-labeled 42-bp single-stranded DNA probe. Following washes, bound DNA was resolved by SDS-PAGE and visualized directly by in-gel Cy5 fluorescence. SPRTN^E112Q^ bound the DNA probe substantially above the GFP-only background (∼ 43% bound), and this binding was reduced by prior treatment of cells with H2O2 to induce cellular PARylation (26.9% bound). DNA binding by the SPRTN^PARless-E112Q^ mutant bound similarly to SPRTN^E112Q^ (47.2% bound). However, unlike SPRTN^E112Q^, SPRTN^PARless-E112Q^ was not reduced by H2O2 treatment (52.4% bound) (**Figure S3A**). Moreover, treatment of the immobilized SPRTN^E112Q^ complexes with PARG restored DNA binding following H2O2 exposure, reversing the inhibitory effect of cellular PARylation (**Figure S3B**). These findings indicate that PARylation of SPRTN directly antagonizes its DNA-binding capacity and that removal of PAR is sufficient to restore SPRTN engagement with ssDNA. To confirm the specificity of this interaction, we performed competition assays using unlabeled ssDNA or RNA. Increasing concentrations of unlabeled ssDNA competed efficiently with Cy5-DNA for binding to GFP-SPRTN^E112Q^, while unlabeled RNA competed only modestly across the same concentration range (**Figure S3C**), indicating that SPRTN preferentially engages single-stranded DNA over RNA.

Having established that PARylation suppresses SPRTN binding to ssDNA and that this effect can be reversed by PARG, we next asked whether PARylation is sufficient to inhibit SPRTN-DNA interactions in a purified system. Catalytically inactive SPRTN^E112Q^ was incubated with recombinant PARP1 and HPF1 in the presence or absence of NAD+, and binding to a FAM-labeled single-stranded DNA substrate was assessed by electrophoretic mobility shift assay (EMSA). In the absence of NAD+, increasing concentrations of SPRTN^E112Q^ generated a concentration-dependent shift of the DNA probe into a slower-migrating SPRTN:DNA complex (**Figure 5E, lanes 1-5**). In contrast, when NAD+ was included to support PARylation, formation of the SPRTN:DNA complex was markedly impaired across the same SPRTN concentration range, and the probe remained predominantly in the unbound form (**Figure 5E, lanes 6-10**). Control reactions confirmed that SPRTN^E112Q^ alone bound DNA, whereas PARP1/HPF1 formed a distinct DNA-bound complex in the absence of NAD+. Notably, this complex was similarly diminished following PARylation (**Figure 5E, lanes 11-15**). Thus, reconstitution of the PARylation reaction with purified components was sufficient to recapitulate the loss of DNA binding observed in cells. Together with the resistance of SPRTN^PARless^ to H2O2-induced inhibition and the restoration of DNA binding by PARG treatment, these findings demonstrate that PARylation directly and reversibly suppresses SPRTN association with DNA.

### SPRTN^PARless^ is hyperactive, drives off-target proteolysis, and accelerates DPC resolution

SPRTN is known to undergo autocatalytic cleavage as a hallmark of its protease activity when over-expressed, generating characteristic lower-molecular-weight fragments. To ask whether loss of PARylation affects SPRTN’s intrinsic protease activity, we compared cleavage fragment abundance among GFP-SPRTN^WT^, GFP-SPRTN^NHD^ (the PAR-binding-deficient NHD mutant), GFP-SPRTN^PARless^, and a GFP-SPRTN^NHD/PARless^ (double mutant). The NHD mutant showed markedly fewer cleavage fragments than wild-type SPRTN, consistent with a loss-of-function phenotype. In contrast, the PARless mutant displayed increased autocleavage relative to wild-type SPRTN (**Figure 6A**). Combining the NHD mutation with the PARless mutations did not restore cleavage, indicating that the increased activity of the PARless mutant requires an intact NHD. Together with the reduced DNA binding of the NHD mutant (**Figure S3A**), these data suggest that NHD-dependent engagement with PAR and chromatin is a prerequisite for SPRTN autoproteolysis, and that PARylation acts downstream of this step to limit it. Consistently, H2O2 treatment reduced the relative abundance of cleavage fragments for both GFP-SPRTN^WT^ and GFP-SPRTN^PARless^, although SPRTN^PARless^ retained a higher relative level of cleavage fragments than SPRTN^WT^ both baseline and after H2O2 treatment (**Figure S4A**). We next compared GFP-SPRTN^WT^, GFP-SPRTN^PARless^, and their respective catalytically dead controls (E112Q), which lack protease activity and therefore define which lower-molecular-weight bands represent genuine autocleavage products rather than nonspecific species. SPRTN^PARless^ showed an increased abundance of cleavage fragments relative to SPRTN^WT^, while both SPRTN^E112Q^ controls showed no cleavage fragments (**Figure 6B**), confirming that loss of PARylation increases SPRTN’s autoproteolytic activity.

**Figure 6:**
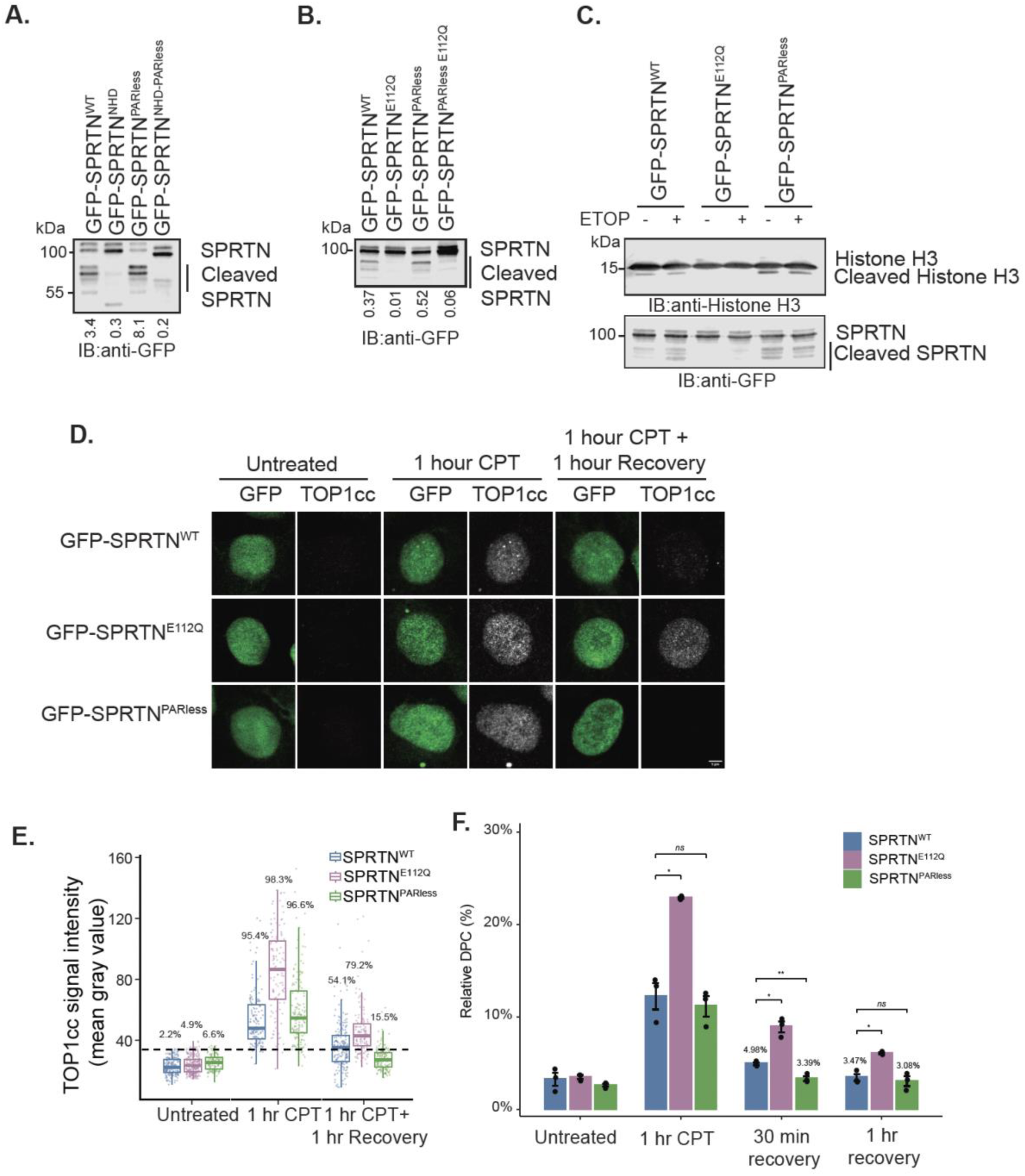
**SPRTN^PARless^ is hyperactive, drives off-target proteolysis, and accelerates DPC resolution A-B**. Whole-cell extracts from HEK293T cells expressing SPRTN variants were analyzed by immunoblotting. The ratio of cleaved fragments to full length SPRTN is indicated below each lane. **A.** GFP-SPRTN^WT^, GFP-SPRTN^NHD^, GFP-SPRTN^PARless^ and GFP-SPRTN^NHD-PARless^. **B.** GFP-SPRTN^WT^, GFP-SPRTN^E112Q^, GFP-SPRTN^PARless^ and GFP-SPRTN^PARless-E112Q^. **C.** HEK293T cells expressing GFP-SPRTN^WT^, GFP-SPRTN^E112Q^ and GFP-SPRTN^PARless^ were subjected to chromatin fractionation. Cells were treated with and without etoposide (ETOP). Only the chromatin-bound fraction is shown. **D.** Representative images from the TOP1cc assay. Dox inducible U2OS Flp-In cells expressing GFP-SPRTN^WT^, GFP-SPRTN^E112Q^ or GFP-SPRTN^PARless^ were either untreated or treated with camptothecin (CPT, 20 μM) for 1 hour followed by 1 hour recovery. Cells were stained with anti-TOP1cc to visualize TOP1cc signal. GFP fluorescence indicates protein expression. **E.** Quantification of the TOP1cc assay. Approximately 150-300 cells were analyzed for each condition. The x-axis represents experimental conditions, and the y-axis represents TOP1cc signal intensity (mean gray value). The dashed line represents the signal threshold. Cells above the threshold are considered TOP1cc positive, and percentages are indicated for each condition. **F.** ARK assay. U2OS Flp-In cells expressing GFP-SPRTN^WT^, GFP-SPRTN^E112Q^ and GFP-SPRTN^PARless^ were either untreated or treated with camptothecin (CPT, 20 μM) for 1 hour followed by 30 minutes or 1 hour recovery. The x-axis represents treatment conditions, and the y-axis represents the ratio of DPC DNA to free DNA. Statistical comparisons for recovery conditions were performed using two-tailed t-tests on raw values: 1 hour CPT, WT vs E112Q, p = 0.02; WT vs PARless, p = 0.157; 30 min recovery, WT vs E112Q, p = 0.031; WT vs PARless, p = 0.005; 1 hour recovery, WT vs E112Q, p = 0.013; WT vs PARless, p = 0.567. * p < 0.05, ** p < 0.01.

Considering that SPRTN’s protease domain shows only limited substrate specificity, we asked whether SPRTN^PARless^ hyperactivity comes at the cost of proteolysis of chromatin proteins not directly involved in DPC repair. We expressed GFP-SPRTN^WT^, GFP-SPRTN^E112Q^, or GFP-SPRTN^PARless^ in cells, treated with or without the topoisomerase II poison etoposide, and immunoblotted for histone H3. A cleaved histone H3 species was detectable in cells expressing SPRTN^WT^ independent of etoposide treatment, indicating that SPRTN overexpression alone is sufficient to drive some degree of histone H3 proteolysis. This cleaved species was absent in cells expressing the catalytically inactive SPRTN^E112Q^ mutant, confirming that it depends on SPRTN’s protease activity rather than another etoposide-or overexpression-associated process^5^. Notably, cells expressing SPRTN^PARless^ showed more cleaved histone H3 than those expressing SPRTN^WT^ (**Figure 6C**). While SPRTN^WT^ itself produces some histone H3 cleavage, the further increase seen with SPRTN^PARless^ is consistent with a model in which PARylation normally limits the extent of SPRTN-mediated proteolysis of chromatin-associated proteins beyond its intended DPC substrates.

To determine whether this increased proteolytic activity corresponds to enhanced DNA-protein crosslink (DPC) resolution in cells, we used topoisomerase I cleavage complexes (TOP1cc) as a defined DPC substrate. Cells expressing GFP-SPRTN^WT^, GFP-SPRTN^E112Q^, or GFP-SPRTN^PARless^ (**Figure S4B**) were treated with 20 µM camptothecin (CPT) for 1 hour to induce TOP1cc formation, then either fixed immediately or allowed to recover for 1 hour after CPT washout, and the percentage of cells positive for TOP1cc staining was quantified by immunofluorescence (**Figure 6D**). Nearly all cells showed TOP1cc-positive nuclei immediately after CPT treatment across all three genotypes (95.4%, 98.3%, and 96.6% of cells for WT, E112Q, and PARless, respectively), confirming comparable initial damage burden. As expected, the SPRTN^E112Q^ mutant showed the least resolution of TOP1cc after 1 hour of recovery (79.2% of cells remained positive). Cells expressing SPRTN^WT^ showed substantially more resolution (54.1% remaining positive), and cells expressing SPRTN^PARless^ resolved TOP1cc most efficiently of the three genotypes, with only 15.5% of the cells remaining TOP1cc-positive after recovery (**Figure 6D and 6E**).

To ask whether faster clearance extends to total CPT-induced DPCs, using an orthogonal, image-independent approach, we used a modified RADAR-based assay (ARK) that biochemically quantifies the fraction of genomic DNA that remains covalently protein-bound relative to total DNA^35^. Consistent with the immunofluorescence data, the ratio of protein-bound to total DNA increased across all three genotypes after 1 hour of CPT treatment. During recovery, cells expressing SPRTN^E112Q^ retained significantly more protein-bound DNA than SPRTN^WT^ at both 30 minutes and 1 hour of recovery, consistent with its catalytic dead status. At 30 minutes of recovery, cells expressing SPRTN^PARless^ showed significantly less protein-bound DNA than SPRTN^WT^; by 1 hour of recovery, protein-bound DNA levels in PARless and wildtype cells were no longer significantly different, as both genotypes had resolved the majority of crosslinks by this time point (**Figure 6F**). Together, these data show that SPRTN^PARless^ accelerates the clearance of TOP1cc, as measured by immunofluorescence and of total CPT-induced DPCs as measured by the ARK assay, with the clearest difference at early recovery time points (**Figure 6D-F**). With the TOP1cc immunofluorescence data, these results demonstrate that PARylation normally restrains SPRTN’s DPC-resolving activity, and that SPRTN^PARless^ resolves TOP1cc more efficiently than wild-type SPRTN, particularly at earlier recovery time points. Taken together, our data support a model (**Figure 7**) in which SPRTN accumulation at DNA-protein crosslinks requires PARP1/2 activity, but SPRTN’s own subsequent PARylation, which depends on its NHD, restrains how long and how well SPRTN remains engaged with damaged chromatin. This PARylation event directly inhibits SPRTN’s DNA-binding and proteolytic activity. Removal of PAR relieves this inhibition, permitting continued or repeated SPRTN engagement with the DPC substrate. Loss of the PARylated residues (SPRTN^PARless^) uncouples SPRTN from this regulatory feedback, yielding a hyperactive protease that exhibits enhanced proteolysis chromatin-associated proteins and resolved DPCs, including TOP1cc, more rapidly than wild-type SPRTN.

**Figure 7.**
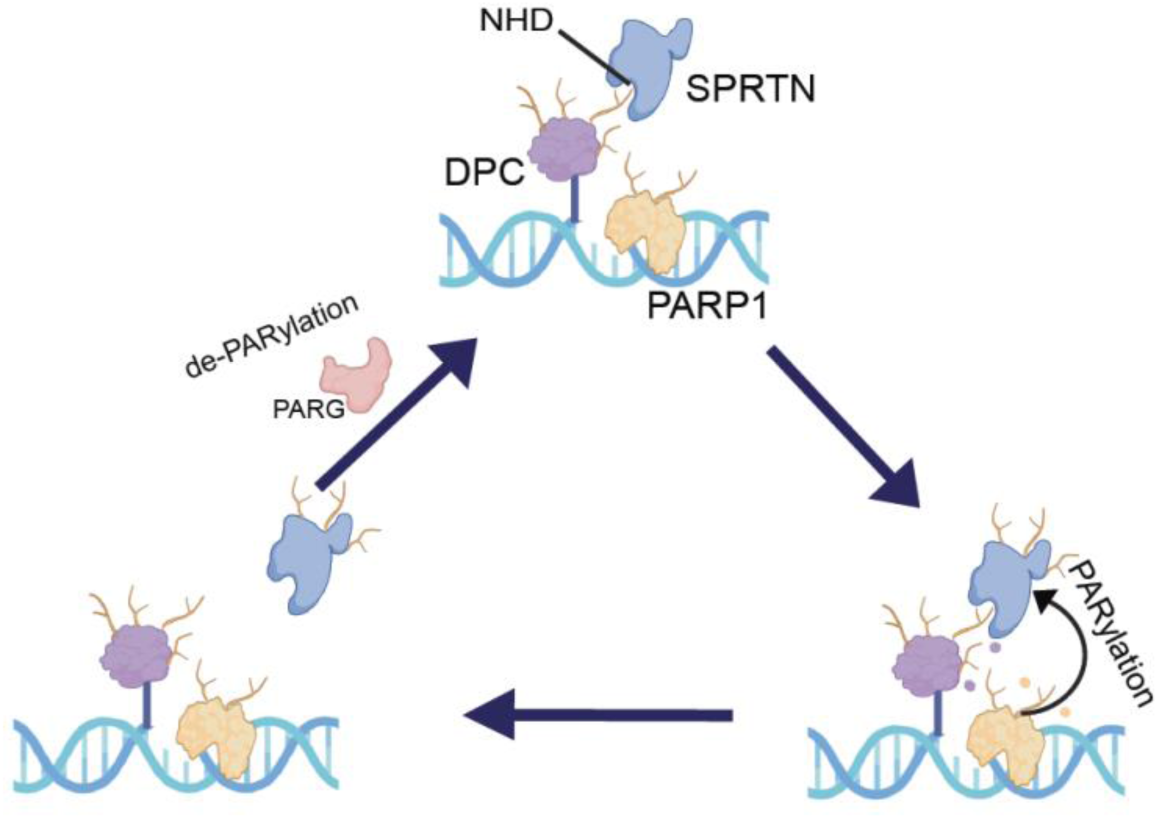
PARylation restrains SPRTN DNA interaction. Model of dynamic PARylation-dePARylation cycles regulating SPRTN DNA engagement and proteolytic activity during DNA damage.

## DISCUSSION

Here we show that SPRTN is a direct substrate of PARP1-HPF1 and that this modification acts as a brake on its activity. PARP1/2 activity is required for SPRTN to accumulate at sites of damage, consistent with the identification of the SPRTN NHD as a PAR reader^25^. Our work adds a second, downstream function for PAR: once SPRTN is engaged, PARP1-dependent modification of its C-terminal region limits its DNA binding, its accumulation on damaged chromatin, and its proteolytic output. We propose that SPRTN is regulated by a recruit-then-restrain cycle in which the same signal that brings the protease to a lesion also limits how long and how actively it works there. This resembles PARP1 itself, whose auto-modification promotes its release from DNA, and it suggests that PAR serves as both a recruitment cue and a timer for DPC proteolysis.

The concentration of PARylation sites in the C-terminus, around the SHP box, PIP box, and UBZ domain, places the modification in a region that connects SPRTN to its partners at the replication fork and to ubiquitinated substrates. The negative charge of PAR chains in this region is well suited to antagonize DNA engagement, as shown for other PARylated DNA-binding proteins, and we show that PARylation reduces SPRTN-DNA binding in cells and in a reconstituted system. Importantly, this effect is reversible by PARG and absent in the PARless mutant, which binds DNA normally but is insensitive to H2O2-induced PARylation. The inhibition therefore depends on the modification itself and can be reset by dePARylation. Whether the effect of PAR is purely electrostatic or also involves a conformational change is a question for future structural work.

Our use of H2O2 as a PARP1 activator places this regulation in the context of oxidative stress. Oxidative damage generates abundant single-strand breaks and base lesions that strongly activate PARP1 across chromatin, and PAR is therefore a general damage signal that does not by itself identify a DNA-protein crosslink. A protease that reads PAR for recruitment but has limited substrate specificity needs a way to avoid acting wherever PARP1 is active. SPRTN’s own PARylation could provide one: when PARP1 activity is high, SPRTN is recruited but constrained, and in our experiments H2O2 reduced SPRTN accumulation at damage in a manner that depended on both PARP1/2 and the SPRTN PARylation sites (**Figure 5A-B**). We suggest that SPRTN uses PAR as a coarse marker of damage and relies on additional cues for substrate selection. Ubiquitin on the crosslinked protein, which allosterically stimulates SPRTN^17,18^, is a strong candidate. In this coincidence-detection model, PAR recruits SPRTN, ubiquitinated substrate licenses proteolysis, and SPRTN’s PARylation limits the duration of its activity. This framework predicts that oxidatively generated DPCs, such as those formed when repair enzymes become trapped at oxidized abasic sites, or by the reactive aldehydes produced during oxidative stress^36–38^, are processed by SPRTN under the control of this cycle. It also predicts that the PARless mutant would show greater non-specific proteolysis during oxidative stress, an idea supported by its increased autocleavage and histone H3 cleavage (**Figure 6B-C**) that is now ready to be tested more broadly.

The functional consequences of PARylation are evident in DPC repair. SPRTN-PARless resolved TOP1cc more rapidly than wild-type SPRTN by immunofluorescence and cleared total CPT-induced DPCs more rapidly by the ARK assay (**Figure 6D-F**), indicating that PARylation sets the pace of SPRTN-dependent repair. A cycle of PARylation and dePARylation could thus allow SPRTN to engage in a crosslink, act, and release, which fits the transient nature of PAR at sites of damage. PARG removes PAR chains while leaving the terminal serine-linked ADP-ribose, which ARH3 removes. That PARG alone restores DNA binding suggests the chain is the inhibitory element, and identifying the physiological dePARylase(s) for SPRTN, and how their activity is coordinated with lesion resolution, is an interesting direction.

Finally, these findings add to a growing picture of SPRTN regulation by phosphorylation, ubiquitination, acetylation, and now ADP-ribosylation, with the C-terminal region as a hub for signal integration. Because both loss of SPRTN function and its dysregulation threaten genome stability, as illustrated by Ruijs-Aalfs syndrome and the toxicity of excess proteolysis, mechanisms that tune SPRTN activity are likely to matter for aging and cancer. Our model gives a framework for studying how SPRTN is controlled in contexts of high PARP activity, such as oxidative stress and treatment with PARP inhibitors, which may alter SPRTN activity and the repair of trapped PARP1 and TOP1 complexes. Further work is needed to define the interplay between the PAR-binding NHD and the C-terminal PARylation sites, to determine whether the PARless protein affects genome stability or sensitivity to DPC-inducing agents, and to extend these findings to endogenous SPRTN.

## ACKNOWLEDGEMENTS

We thank Nicholas Lakin (University of Oxford) for the U2OS PARP1/2 double knockout cell line. This work was supported by the National Science Foundation CAREER award MCB-2146384.

---

**Figure S1.**
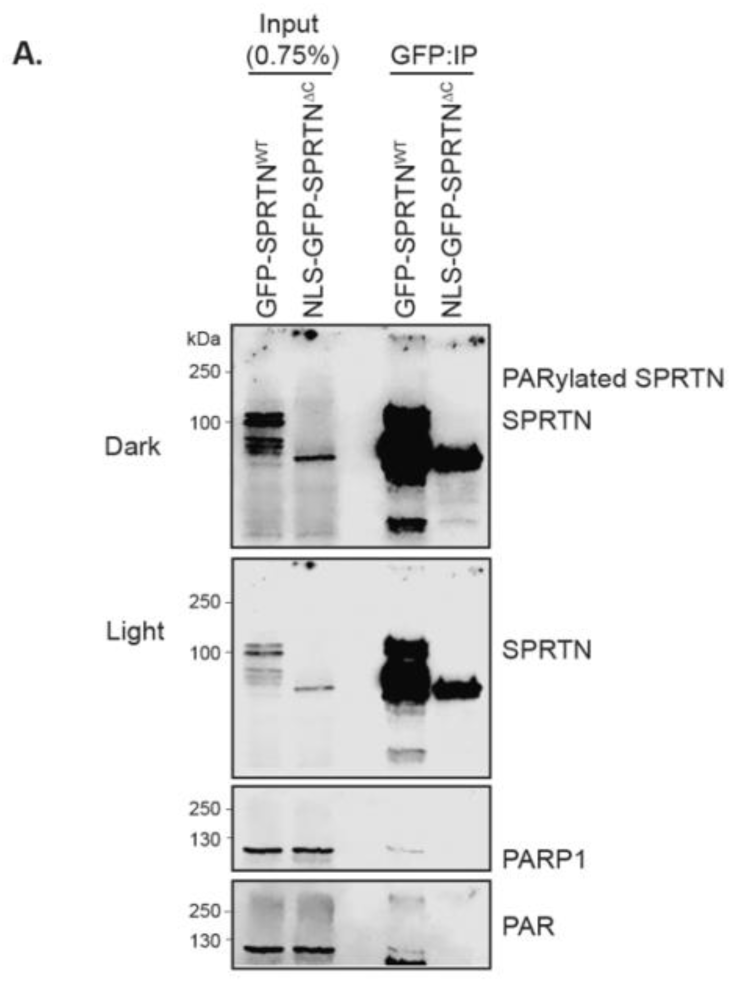
GFP tagged SPRTN^WT^ and SPRTN^ΔC^ were overexpressed and immunoprecipitated from HEK293T cells. NLS-GFP-SPRTN^ΔC^ lacks high mw PARylated SPRTN species.

**Figure S2.**
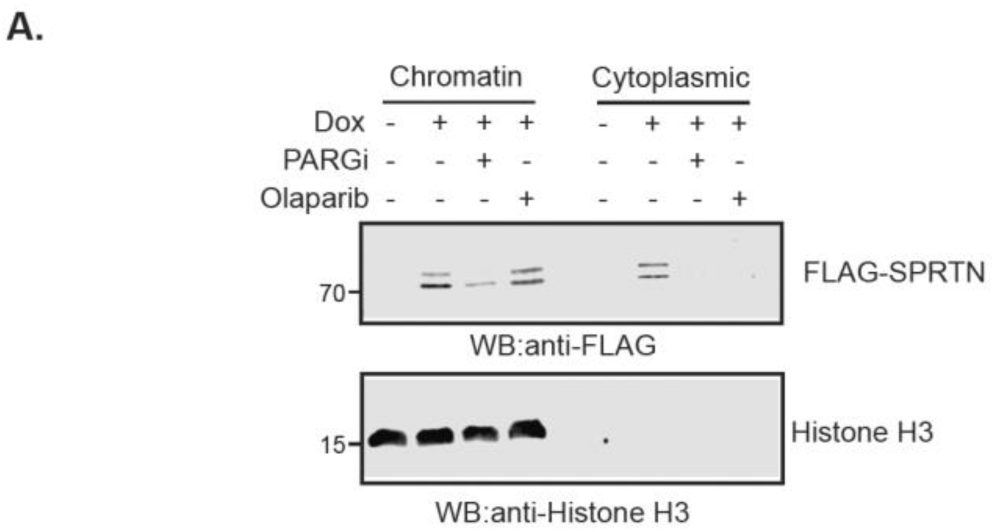
Chromatin fractionation was performed in U2OS Flp-In cells expressing FLAG-SPRTN^WT^ after treatment with PARG inhibitor (10 μM, 12 hours) or Olaparib (20 μM, 12 hours).

**Figure S3.**
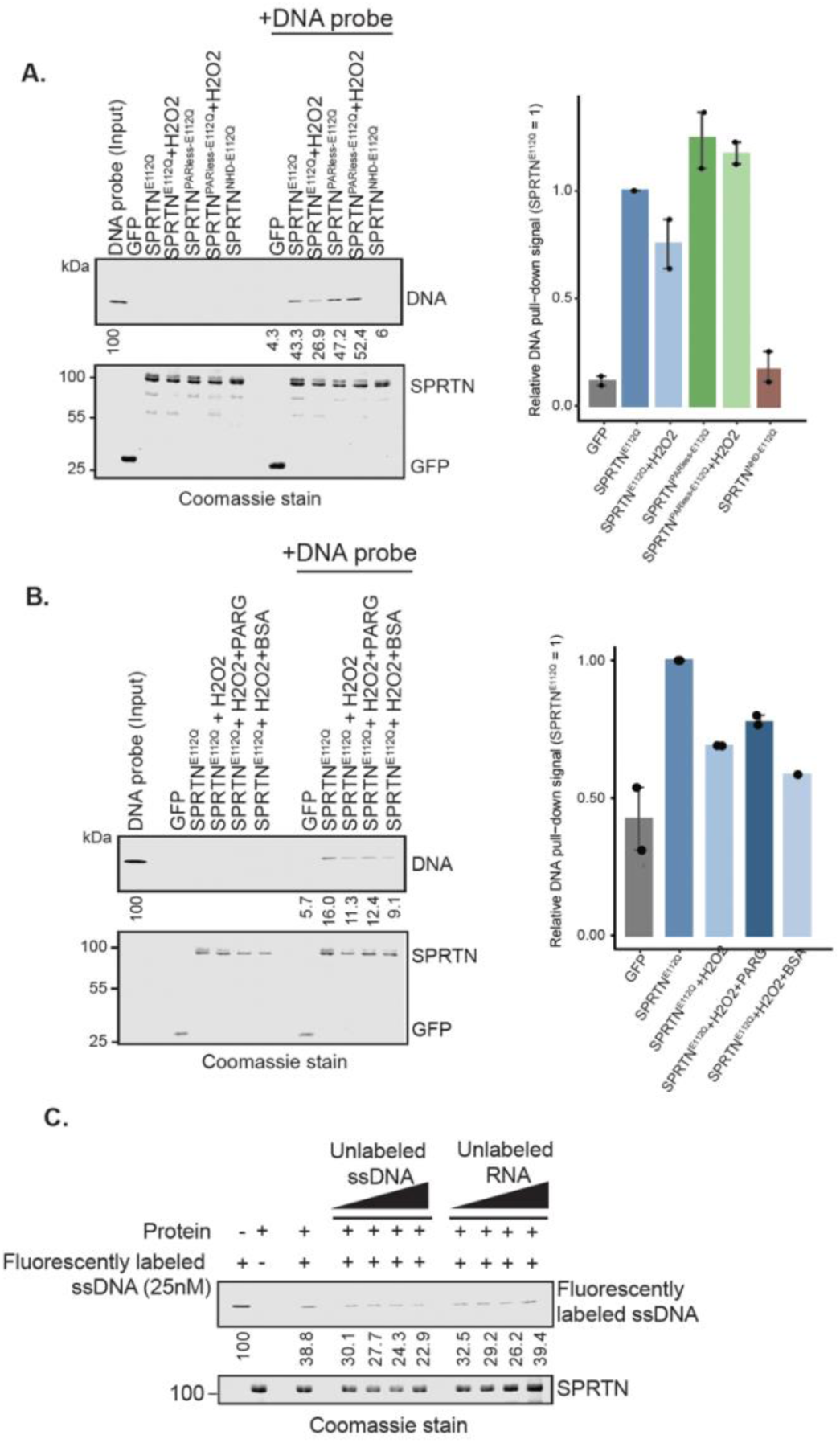
(**A-B**) Immunoprecipitated GFP-SPRTN variants from HEK293T cells treated with H2O2 (5 mM, 15 minutes) were incubated with Cy5-labeled 42-bp single-stranded DNA probe. The percentage of DNA binding relative to input is indicated below for each lane. Coomassie staining shows protein levels. Quantification of relative DNA pull-down is shown on the right. Values are normalized to SPRTN^E112Q^ (set to 1). N=2 **A.** Immunoprecipitated GFP-SPRTN^E112Q^ and GFP-SPRTN^PARless-E112Q^ from both untreated and H2O2 treated cells were incubated with Cy5-labeled 42-bp single-stranded DNA probe. GFP-SPRTN^NHD-E112Q^ served as DNA binding control. **B.** GFP-SPRTN^E112Q^ immunoprecipitated from H2O2 treated cells was subjected to PARG treatment (1 hour) prior to incubation with Cy5-labeled ssDNA. BSA served as a treatment control. **C.** Immunoprecipitated GFP-SPRTN^E112Q^ was incubated with Cy5-labeled ssDNA in the presence of increasing concentrations (12.5, 25, 50 and 100 nM) of unlabeled ssDNA or RNA as competitors.

**Figure S4.**
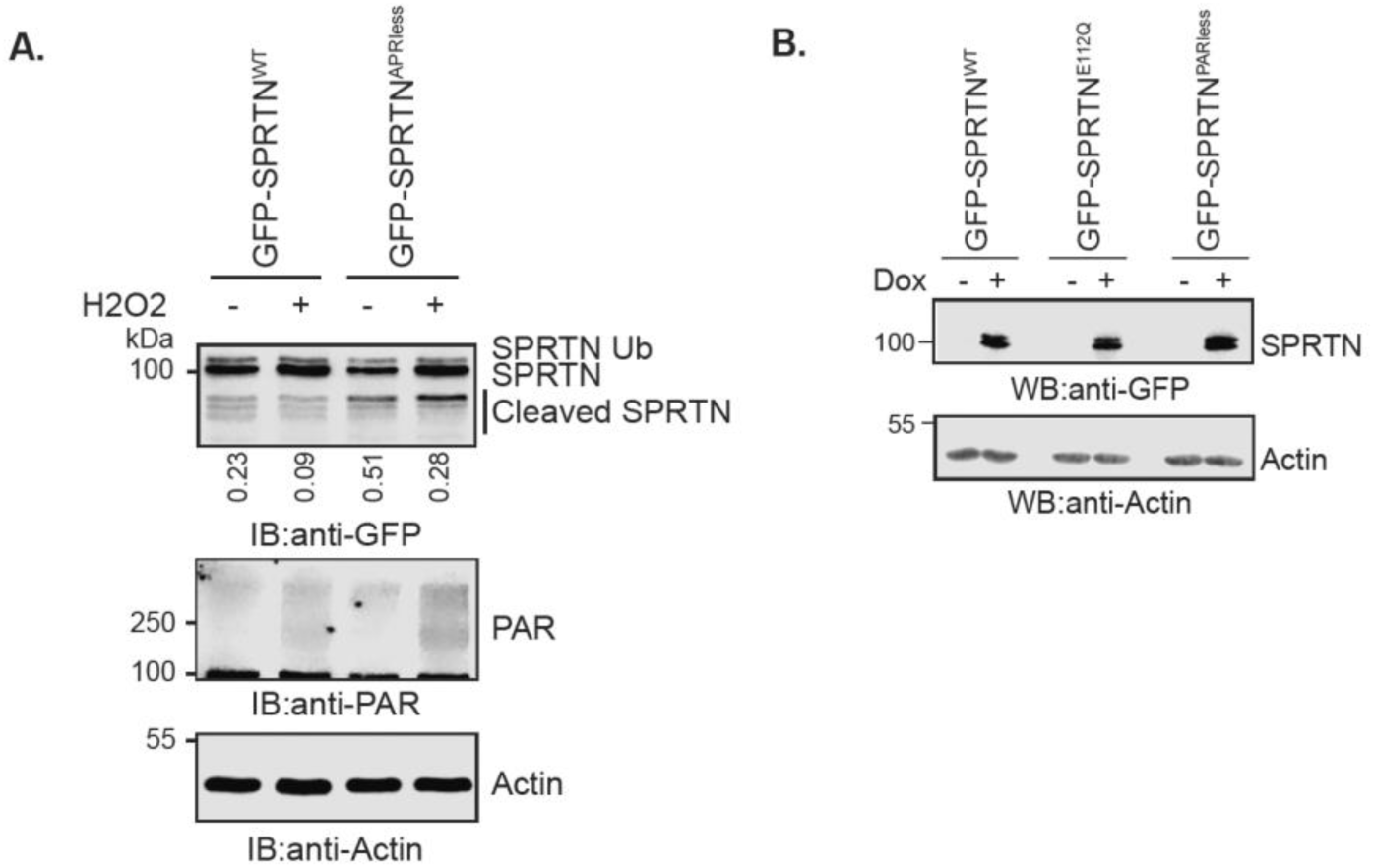
A. Whole-cell extracts from HEK293T cells expressing GFP-SPRTN^WT^ and GFP-SPRTN^PARless^ treated with or without H2O2 (5 mM for 15 minutes). The ratio of SPRTN cleavage fragments to full length SPRTN is indicated below each lane. **B.** Whole-cell extracts from inducible U2OS Flp-In cells expressing GFP-SPRTN^WT^, GFP-SPRTN^E112Q^ and GFP-SPRTN^PARless^ were analyzed by anti-GFP immunoblotting. Anti-actin immunoblotting served as a loading control.

## MATERIALS AND METHODS

### Cell Culture

HEK293T and U2OS Flp-In cells were cultured in Dulbecco’s Modified Eagle Medium (DMEM) supplemented with 10% fetal bovine serum (FBS), penicillin and streptomycin. Transient transfections were performed using either TurboFect or PolyJet following manufacturer’s guidelines and cells were harvested after 24 hours of post-transfection. Treatment with Doxycycline (1 μg/mL, overnight), PARGi (10 μM, overnight), Olaparib (20 μM, overnight), H2O2 (5 mM, 15 min), Etoposide (5 μM, 1hr), or CPT (20 μM, 1 hour) were performed as indicated in each experiment.

### Cloning and Site-Directed Mutagenesis

SPRTN variants were cloned into pEGFP to generate N-terminally tagged GFP-SPRTN^WT^, GFP-SPRTN^PARless^, GFP-SPRTN^NHD^ and GFP-SPRTN^NHD-PARless^ constructs.

The Q5® Site-Directed Mutagenesis Kit (New England Biolabs) was used to create GFP-SPRTN^E112Q^, GFP-SPRTN^E112Q^, GFP-SPRTN^NHD-E112Q^ and GFP-SPRTN^PARless-E112Q^ in the same vector backbone.

To generate a doxycycline inducible system, SPRTN variants were also cloned in pcDNA5 to produce N-terminally tagged GFP-SPRTN^WT^, GFP-SPRTN^E112Q^ and GFP-SPRTN^PARless^ constructs. These constructs were made shRNA resistant by site-directed mutagenesis. shRNA control and shRNA-SPRTN sequences were cloned in mCherry-pGIPZ. For protein expression and purification, PARG and USP2 were cloned in pGex6p1 vector. All plasmid sequences were verified using the Illumina-based NGS platform provided by Plasmidsaurus.

### Cell Line Generation

shRNA Control and shRNA-SPRTN cell lines were generated in U2OS Flp-In cells using lentiviral transduction. To produce lentivirus, HEK293T cells were transfected with lentiviral packaging vectors psPAX2 and VSV-G (pMD2.G) along with the transfer plasmid using Lipofectamine, following the manufacturer’s protocol. Lentiviral supernatant was collected 48 hours post-transfection and centrifuged at 4200 rpm for 15 min to remove cell debris. Recipient U2OS Flp-In cells were transduced with the lentiviral supernatant for 24 hours. Cells were allowed to recover from transduction before initiating antibiotic selection. The knockdown efficiency of the resulting shRNA Control and shRNA-SPRTN U2OS Flp-In cell lines was confirmed by RNA sequencing from Plasmidsaurus, followed by differential gene expression analysis.

The shRNA-SPRTN U2OS Flp-In cells were subsequently used to generate doxycycline-inducible stable cell lines expressing GFP-SPRTN^WT^, GFP-SPRTN^E112Q^, GFP-SPRTN^PARless^. For Flp recombination, cells were transfected with pOG44 (900 ng) and the transfer plasmid (100 ng) using Lipofectamine, following the manufacturer’s protocol. Cells were allowed to recover prior to antibiotic selection. Stable integration and protein expression were confirmed by Western blot analysis.

### Immunoblotting (IB) and Immunoprecipitation (IP)

For immunoblotting (IB), lysis was performed directly in 1X Laemmli buffer (0.031M Tris pH 6.8, 1% SDS, 2.5% β-mercaptoethanol, 5% glycerol, 0.125 mg/mL bromophenol blue), followed by boiling (90°C for 5 min) and sonication.

For immunoprecipitation (IP), cell pellets were lysed in buffer containing 25 mM Tris-HCl (pH 7.5), 150 mM NaCl, 0.1% NP-40, 10% glycerol, 2 mM EDTA, 1 mM PMSF, 1 mM DTT, protease inhibitor, 10 μM PARG inhibitor, Benzonase and 2 mM MgCl2 on ice for 15 min. Lysates were sonicated on ice and clarified by centrifugation at 15000 rpm for 15 min at 4°C. Clarified lysates were incubated with the equilibrated GFP-Trap beads for 2 hours at 4°C on a rotating platform to immunoprecipitate GFP-tagged proteins. Beads were washed four times with wash buffer (25 mM Tris HCl pH 7.5, 150 mM NaCl, 0.1% NP-40, 10% glycerol, 2 mM EDTA, 1 mM PMSF, 1 mM DTT, 10 μM PARG inhibitor).

Immunoprecipitated GFP tagged proteins were treated with PARG and/or 2 μM USP2 (pre-activated in 25 mM Tris HCl pH 7.5, 150 mM NaCl, 10 mM DTT on ice for 10-15 min) at 37°C for 30 min or 1 hour as indicated. Deubiquitination reaction was performed in 50 mM Tris HCl pH 7.5, 50 mM NaCl and 5 mM DTT. Samples were then prepared for immunoblotting by adding 2X Laemmli buffer. Protein samples were resolved by SDS-PAGE and transferred onto nitrocellulose membranes for immunoblotting. Membranes were blocked with 5% milk in TBS, followed by primary antibody incubation overnight at 4°C and secondary antibody incubation for 1 hour at room temperature. Membranes were visualized and quantified using a LI-COR imaging system.

The following antibodies were used for immunoblotting: anti-GFP (1:1000), anti-PAR (1:5000), anti-PARP1 (1:1000), anti-Flag (1:20,000), anti-Histone H3 (1:1000), anti-SPRTN (1:10), anti-Actin (1:1000) and anti-ubiquitin (P4D1) (1:1000).

### DNA Pull-Down Assay

DNA pull-down assays were performed using immunoprecipitated GFP-tagged proteins immobilized on GFP-Trap beads. Beads were washed five times with high salt wash buffer (25 mM Tris HCl pH 7.5, 400 mM NaCl, 0.1% NP-40). DNA-binding reactions were performed by adding 25 nM fluorescently labeled ssDNA (42 nt) to the IP material in 1X binding buffer (25 mM Tris HCl pH 7.5, 100 mM NaCl, 1 mM DTT), followed by incubation at 37°C for 1 hour with shaking.

For competition assay, 12.5 nM, 25 nM, 50 nM and 100 nM unlabeled ssDNA or RNA were added as indicated. After incubation, beads were washed five times with binding buffer and samples were prepared in Laemmli buffer prior to SDS-PAGE. Fluorescent DNA was visualized and quantified using a LI-COR imaging system, and protein levels were confirmed by Coomassie staining.

### Biolayer Interferometry (BLI)

HEK293T cells overexpressing GFP-SPRTN^E112Q^ were lysed in 25 mM Tris-HCl (pH 7.5), 150 mM NaCl and 0.1% NP-40 on ice for 15 minutes. Lysates were sonicated and clarified at 15000 rpm for 15 minutes at 4 °C. Whole-cell lysates and biotinylated 20-mer ssDNA were diluted in kinetic buffer (25 mM Tris-HCl pH 7.5, 150 mM NaCl, 0.02% Tween, 0.05% BSA) for the assay. DNA binding kinetics were measured using an Octet R2 biolayer interferometry (BLI) system (Sartorius). Streptavidin biosensors were incubated with 500 nM biotinylated 20-mer ssDNA to load DNA onto the sensor. Non-transfected whole-cell lysate was used as a baseline before the biosensors were exposed to samples to measure binding kinetics. Association and dissociation were monitored for 10 minutes and 3 minutes, respectively. Quantification of data was done in R (RStudio).

### Protein Expression and Purification in *E. coli*

*E. coli* Rosetta (DE3) cells were used to express GST-PARP1 and GST-PARG. Bacterial cultures were grown in LB medium supplemented with 100 μg/mL ampicillin. Protein expression was induced with IPTG (0.5 mM, overnight) once the culture reached an OD_600_ of 0.6 at 25°C. Cells were harvested by centrifugation, washed with PBS and stored at - 80°C until use. Cell pellets were lysed in buffer containing PBS, 380 mM NaCl, 1 mM DTT, protease inhibitor and lysozyme (1 mg/mL) on ice for 30 minutes. Lysates were sonicated and clarified before Glutathione Sepharose beads were added to capture GST tagged proteins. GST-tagged proteins bound to the beads were washed five times with wash buffer (PBS, 380 mM NaCl). PreScission protease (3C) was then used to cleave the GST tag from the purified proteins.

### *In Vitro* PARylation Assay

*In Vitro* poly ADP ribose (PAR) polymer was synthesized to test the activity of purified PARG. DpnI-digested DNA, PARP1, 1 mM NAD+ were incubated in reaction buffer containing 25 mM Tris-HCl pH 7.5 and 100 mM NaCl at 37 °C for 1 hour, followed by the addition of Benzonase (37 °C, 30 min). The reaction was stopped by heat inactivation of PARP1 at 90°C for 5 minutes. Synthesized PAR was then incubated with PARG at 37 °C for 30 minutes to test PARG activity. *In vitro* SPRTN PARylation reactions contained 0.5 μM full-length SPRTN, 1.25 pmol PARP1, 500 μM NAD+, 25 pmol HPF1, and 0.2 μg nucleosomes. Each reaction was brought up to a final volume of 10 μl using reaction buffer (25 mM Tris-HCl pH 8, 100 mM NaCl, and 1 mM DTT). Reactions were incubated at 37°C for 2 hours and stopped by the addition of 4X Laemmli buffer followed by boiling for 3 minutes. Samples were resolved by SDS-PAGE and analyzed by immunoblotting.

### Chromatin Fractionation

Chromatin fractionation was performed as previously described . Cell pellets were resuspended in Buffer A (10 mM HEPES pH 7.9, 10 mM KCl, 1.5 mM MgCl2, 0.34 M sucrose, 10% glycerol, 1 mM DTT, protease inhibitor) supplemented with 0.1% Triton X-100. The cytoplasmic fraction was separated from the nuclear fraction by centrifugation at 1300 x g for 5 minutes at 4 °C. The nuclear pellet was washed three times with Buffer A and lysed in Buffer B (3 mM EDTA, 0.2 mM EGTA, 1 mM DTT, protease inhibitor) for 30 minutes. Centrifugation was performed at 1700 X g for 5 minutes at 4°C to separate the chromatin bound fraction as a pellet. The pellet was washed three times with Buffer B and resuspended in Laemmli buffer. Ten percent of the total chromatin-bound fraction was loaded for immunoblot analysis.

### Immunofluorescence (IF)

TOP1cc assays were performed as previously described . Cells were seeded on sterile coverslips and induced with doxycycline (1 μg/mL, overnight). CPT (20 μM, 1 hour) treatment was performed with either no recovery or 1 hour recovery. Cells were washed once with PBS followed by fixation with 4% PFA in PBS (4 °C, 15 min) and permeabilization with 0.5% SDS in PBS for 5 minutes. Cells were washed five times with wash buffer (PBS containing 0.1% Triton X-100, 0.1% BSA) and blocked in 5% BSA for 1 hour. Cells were then incubated with anti-TOP1cc primary antibody (1:200) in 2.5% BSA in PBS for 1 hour at room temperature. After primary antibody incubation, cells were washed three times with PBS and incubated with Alexa Fluor 647 secondary antibody (1:500) diluted in 1% BSA in PBS for 1 hour at room temperature. Finally, cells were washed three times with PBS and stained with DAPI. Imaging was performed using an Olympus FV1000 confocal microscope with a 40x objective. For each condition, ∼150-300 cells were imaged for quantification. All images were acquired using similar laser power and exposure settings.

### ARK assay

4×10^5^ U2OS Flp-In stable cells expressing GFP-SPRTN^WT^, GFP-SPRTN^E112Q^ and GFP-SPRTN^PARless^ were seeded in triplicate and treated overnight with doxycycline to induce protein expression. Cells were treated with CPT (20 μM, 1 hour) and allowed to recover for either 30 minutes or 1 hour. At the time of collection, cells were washed twice with PBS and lysed in 700 μL pre-warmed M-buffer (5.6 M GTC, 10 mM Tris-HCl pH 6.8, 20 mM EDTA, 4% Triton X-100, 1% Sarkosyl, 1% DTT) for 10 minutes at room temperature. Lysates were triturated 30 times using a 200 μL pipette tip, followed by mixing with an equal volume of cold ethanol (−20 °C) and centrifugation at 20,000 x g for 30 minutes at 4°C. Pellets were washed three times with Tris-HCl buffer (20 mM Tris-HCl pH 6.8, 150 mM NaCl, 50% ethanol) and subsequently resuspended in 500μL prewarmed SDS buffer (1% SDS, 20 mM Tris-HCl pH 7.5) and incubated at 42°C for 6 minutes. DNA was sheared using a 27-gauge needle, after which 500 μL of KCl buffer (200 mM KCl, 20 mM Tris-HCl pH 7.5) was added and the samples were incubated on ice for 6 minutes. Samples were centrifuged at 20,000 x g for 5 minutes at 4 °C, and the supernatant was collected for free DNA measurement. The precipitate was washed twice by resuspending in wash buffer (100 mM KCL, 20 mM Tris-HCl pH 7.5), incubating at 55°C for 10 minutes, followed by incubation on ice for 4 minutes and centrifugation at 20,000 X g for 5 minutes at 4 °C. Supernatants from each wash were pooled with the previously collected fraction for free DNA measurement. The remaining precipitate was resuspended in 1 mL Proteinase K Buffer (100 mM KCl, 20 mM Tris-HCl pH 7.5, 10 mM EDTA, 0.2 mg/mL Proteinase K) and incubated 55 °C for 45 minutes. Samples were then chilled on ice for 6 minutes and centrifuged at 20,000 x g for 10 minutes in 4 °C. The supernatant was collected and DNA was quantified using a Qubit fluorometer to measure free DNA and protein-bound DNA. Data were reported as protein bound-DNA/ (protein-bound DNA + free DNA) and normalized to untreated samples.

### UV Laser Micro-irradiation (UV-IR)

U2OS cells were seeded in glass-bottom Lab-Tek dishes and transfected with constructs encoding GFP-SPRTN^WT^ or GFP-SPRTN^PARless^. Cells were treated with BrdU (20 μM) overnight prior to micro-irradiation. Localized DNA damage was induced using a 405-nm UV laser at 100% laser intensity on the NSPARC Nikon system. Damage was generated along user-defined lines within individual nuclei, and images were acquired automatically every 30 s for up to 5 min following micro-irradiation. Approximately 20–30 cells were analyzed per condition, and GFP fluorescence intensity at the irradiated region was quantified over time. For experiments examining the effect of PARP activation on SPRTN recruitment, cells were treated with H2O2 (5 mM) prior to UV laser micro-irradiation.

### Electrophoretic Mobility Shift Assay (EMSA)

DNA-binding activity of purified SPRTN was assessed by electrophoretic mobility shift assay (EMSA). A 42-bp single-stranded DNA probe (oJLO644) labeled at the 5′ end with 6-FAM was used at a final concentration of 2.5 nM. Purified 6×His-SPRTN^E112Q^ was incubated at the indicated concentrations (0–2.0 μM) with the DNA probe in binding buffer containing 20 mM HEPES-KOH (pH 8.0), 80 mM KCl, 1 mM DTT, and 0.1% BSA. Binding reactions were prepared to a final volume of 20 μL by combining binding buffer and protein before addition of the DNA probe and were incubated for 30 min at room temperature. Following incubation, 5 μL of 4× native loading buffer (40% glycerol, 0.02% bromophenol blue) was added to each reaction. Samples were resolved on a 6% native polyacrylamide gel prepared in 0.5× TBE. Gels were pre-electrophoresed for 20 min in chilled 0.5× TBE and subsequently run at 100 V for 1.5 h with chilled 0.5× TBE. Following electrophoresis, the gel was imaged using a Typhoon fluorescence imaging system to detect the 6-FAM-labeled DNA probe.

### Image Quantification and Statistical Analysis

All confocal images were analyzed using ImageJ/ Fiji. Signal intensity was quantified as mean grey value after background subtraction. For micro-irradiation experiments, time-course data were presented as mean ± SEM for each time point from ∼20-30 cells per condition. For the TOP1cc assay, thresholding was performed after background subtraction. To identify TOP1cc-positive cells, a cut-off threshold of signal intensity was determined using the mean intensity + 2 SD (standard deviation) of untreated WT samples. Signal intensity above this threshold was classified as TOP1cc-positive. The same threshold was applied uniformly across all samples and conditions. For ARK assay, statistical analysis was performed using unpaired two tailed Student’s t-test, as indicated in the figure legends. Data presented as mean ± SEM, and a p-value < 0.05 was considered statistically significant. Data were exported and plotted in R (RStudio).

---

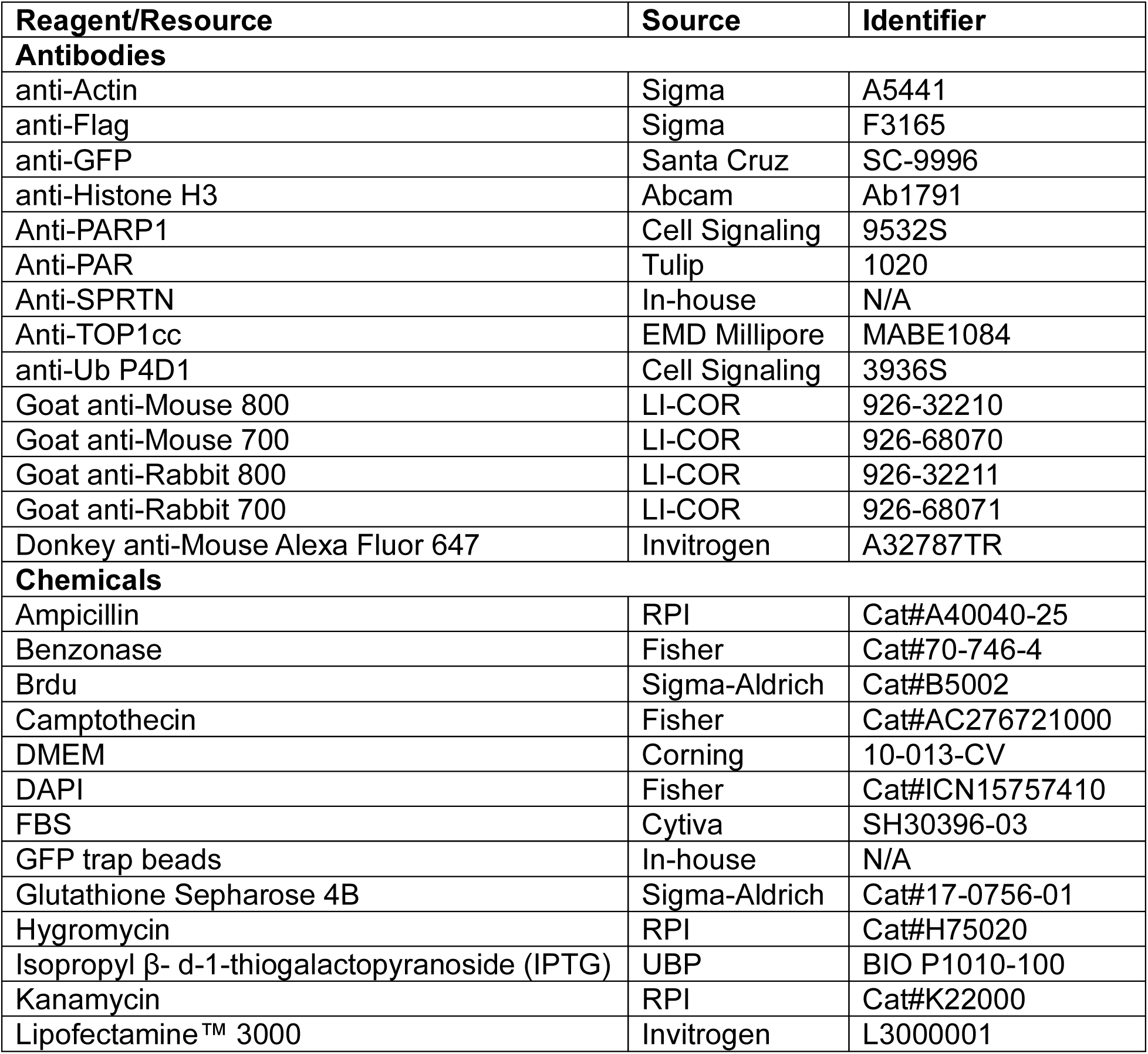
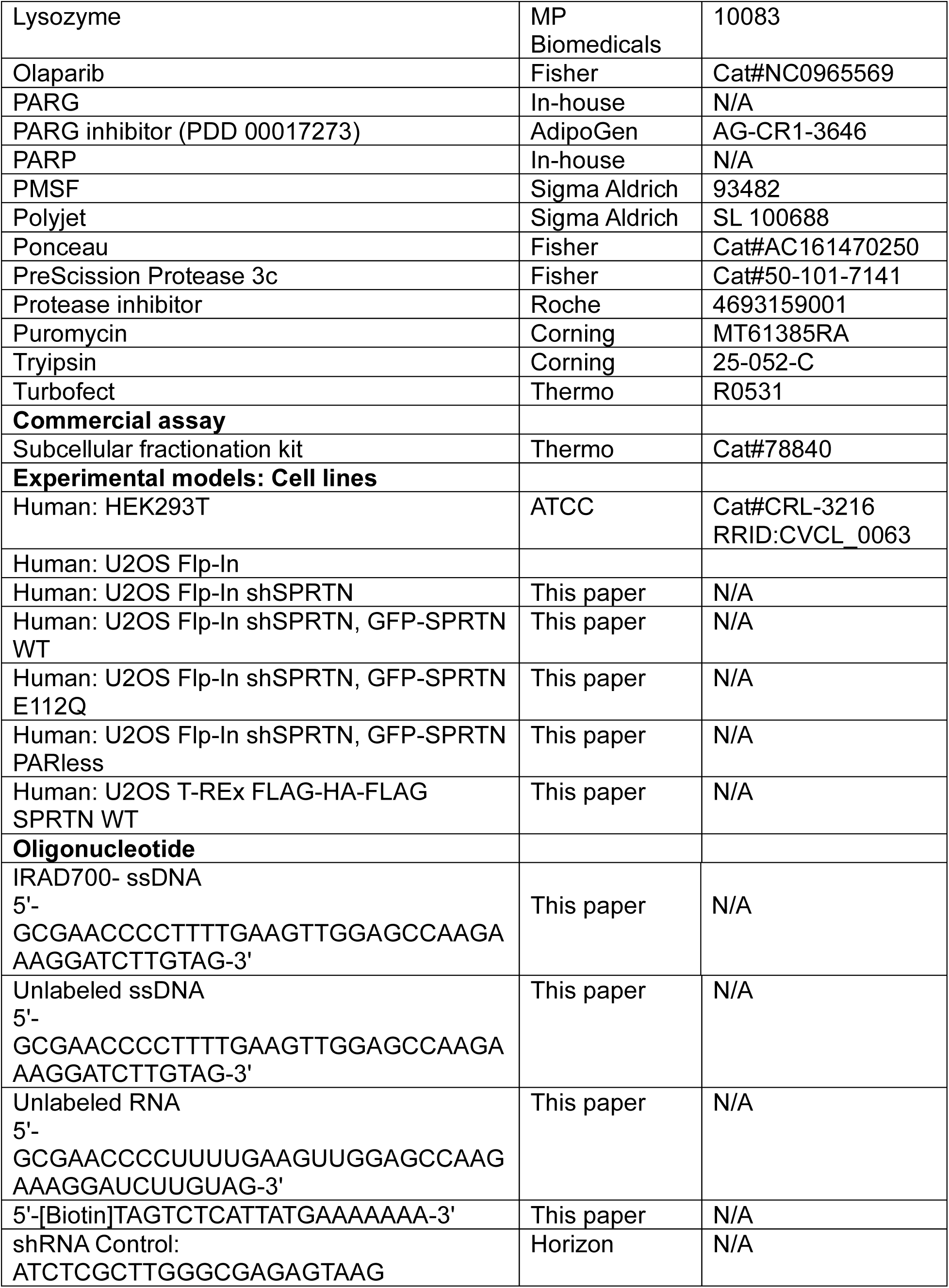
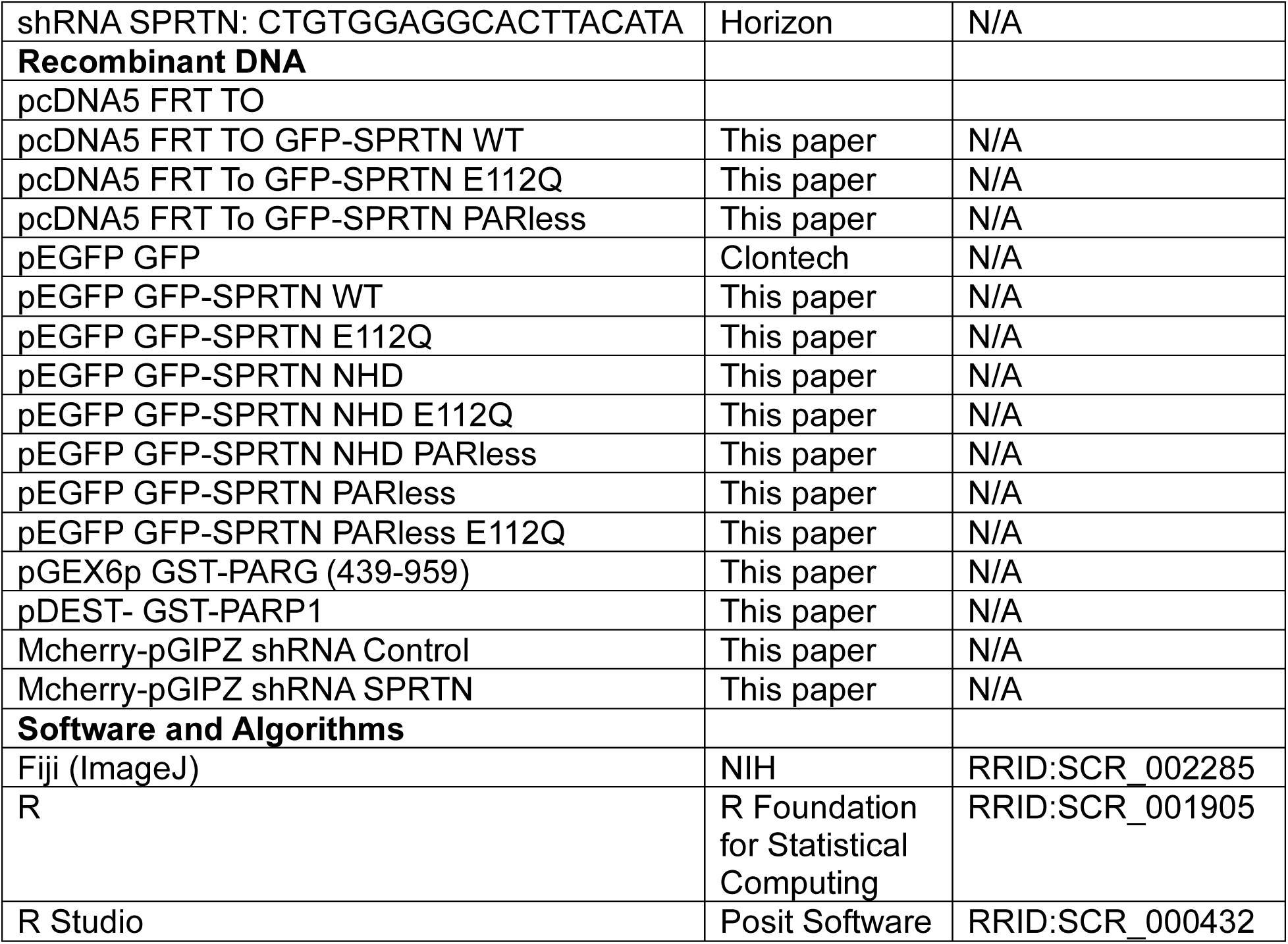

## Notes

### Competing Interest Statement

The authors have declared no competing interest.

